# Zeaxanthin-dependent rapid energy quenching uncovers complexity of PsbS protonation

**DOI:** 10.64898/2026.11.21.768332

**Authors:** Raquel Ponce, Ella Liu, Krishna K. Niyogi, Masakazu Iwai

## Abstract

Rapid responses to changes in light intensity are crucial for photosynthetic organisms to regulate their light-harvesting capacity and optimize growth. PsbS, the S subunit of photosystem II (PSII) located in the thylakoid membrane, senses lumenal pH changes that reflect the balance between the light and carbon reactions of photosynthesis. Upon lumenal acidification caused by absorption of excess light energy, PsbS undergoes protonation of its lumen-exposed domain, triggering energy-dependent quenching (qE), which dissipates excess energy as heat. Despite extensive research, the precise molecular mechanism by which PsbS induces qE remains poorly understood. In this study, rather than relying solely on a PsbS null mutant, we used *Arabidopsis* lines expressing PsbS variants with mutations at the two essential protonatable residues (E122 and E226) in the lumenal domains to investigate the molecular interplay among PsbS, lumenal acidification, and zeaxanthin during qE induction. Our results indicate that this mutant PsbS protein can induce a small qE, the magnitude of which is strongly linked to both zeaxanthin accumulation and lumenal acidification. Furthermore, we found that the loss of the monomeric light-harvesting antenna complex CP29 in this mutant PsbS line reduced the small qE induction by nearly half. Together, our observations suggest that the physical presence of PsbS is essential for qE induction, which is driven by direct or indirect interactions between PsbS and zeaxanthin, potentially mediated by light-harvesting antenna complexes.

## INTRODUCTION

An optimal photosynthetic growth rate in plants and green algae is achieved when light energy is utilized with maximum efficiency and safety. Light absorption by chlorophylls and subsequent excitation energy transfer toward the photosystems leads to charge separation at the reaction centers, which drives photosynthetic electron transport in the chloroplast thylakoid membrane (Blankenship, 2021). Because the rate of electron transport is orders of magnitude slower than the kinetics of excitation energy transfer (Arsenault et al., 2020; Yang et al., 2022; Leonardo et al., 2024), increasing the rate of light absorption can easily saturate the reaction centers, which results in the accumulation of harmful excess energy in chloroplasts (Niyogi, 1999). To minimize the potentially deleterious effects caused by excess energy, the photosynthetic machinery carries out energy dissipation mechanisms, collectively referred to as non-photochemical quenching (NPQ) (Bassi and Dall’Osto, 2021; van Amerongen and Croce, 2025). Since it has been demonstrated that fine-tuning the rate of induction and relaxation of NPQ can enhance photosynthesis and yield in crop plants (Kromdijk et al., 2016; De Souza et al., 2022), there is increasing attention in genetic engineering to modify NPQ activity and improve photosynthetic efficiency in a broader range of crop plants (Long et al., 2022). Nevertheless, the molecular mechanisms of NPQ remain poorly understood, because multiple factors are involved in the induction and relaxation at different timescales.

Among the various known components of NPQ, energy-dependent quenching (qE) is the fastest feedback mechanism, induced within seconds to minutes in response to acidification of the thylakoid lumen under high light (HL) conditions (Murata and Sugahara, 1969; Wraight and Crofts, 1970; Briantais et al., 1979). Earlier studies demonstrate that the extent of NPQ is highly correlated with the accumulation of zeaxanthin (Demmig et al., 1987), which is synthesized from violaxanthin and antheraxanthin by violaxanthin de-epoxidase (VDE) that is activated by the lumenal acidification in the presence of ascorbate (Yamamoto et al., 1962; Yamamoto et al., 1972; Rockholm and Yamamoto, 1996). However, the reconversion of zeaxanthin to violaxanthin by zeaxanthin epoxidase is slower than the de-acidification of the thylakoid lumen, and zeaxanthin can cause energy dissipation through a separate zeaxanthin-dependent quenching mechanism (qZ) that is distinct from qE (Nilkens et al., 2010). In addition, even without zeaxanthin accumulation, a rapid, though smaller, NPQ induction still occurs immediately after light exposure, primarily due to qE triggered by light-dependent lumenal acidification (Niyogi et al., 1998).

About 30 years after the initial detailed studies on qE, the *Arabidopsis npq4* mutant, lacking photosystem II (PSII) subunit S (PsbS), was isolated, which does not exhibit rapid NPQ induction (Li et al., 2000). Although zeaxanthin accumulation in the *npq4* mutant in HL occurs similarly to the wild type (WT), its NPQ capacity is significantly impaired, suggesting that PsbS is required for qE (Li et al., 2000). In addition, the *Arabidopsis* L17 mutant, in which PsbS is overexpressed, shows an enhanced NPQ capacity, indicating that the level of qE is correlated with PsbS protein abundance (Li et al., 2002a). Furthermore, it has been demonstrated that two lumen-exposed glutamate residues of PsbS, namely E122 and E226, are protonated upon lumenal acidification in HL and are responsible for the rapid NPQ induction (Li et al., 2004). Based on computational and spectroscopic analyses, the protonation of E122 and E226 induces conformational changes around these lumen-exposed domains, which potentially affect protein– protein interactions necessary for the rapid NPQ induction (Liguori et al., 2019; Krishnan-Schmieden et al., 2021; Chiariello et al., 2023). As PsbS was originally found to be co-immunoprecipitated with the oxygen-evolving complex of PSII (Ljungberg et al., 1984), several lines of evidence suggest that PsbS is located proximal to PSII and its light-harvesting complex (LHCII) proteins (Correa-Galvis et al., 2016; Sacharz et al., 2017; Daskalakis, 2018; Frances et al., 2025). However, no direct evidence has pinpointed a specific location of PsbS (Teardo et al., 2007; Caffarri et al., 2009), leaving the exact molecular mechanism by which PsbS protonation triggers qE unclear.

PsbS is a member of the light-harvesting complex (LHC) protein superfamily, as it contains transmembrane (TM) helices with the LHC motif (E-X-X-X-X-R-X-A-M) (Jansson, 1999). Though most LHC proteins have three TM helices, PsbS contains four TM helices, which are hypothesized to have arisen through internal gene duplication of a two-helix protein (Kim et al., 1992; Wedell et al., 1992). The crystal structure of PsbS reveals its distinct arrangement of these four TM helices (Fan et al., 2015). TM1 and TM3 form the core structure with twofold symmetry, spanning the membrane plane diagonally, which is similar to the helices A and B in LHCII proteins (Liu et al., 2004; Standfuss et al., 2005). On the other hand, TM2 and TM4 flank the groove of the core structure, creating a more compact conformation than LHCII and lacking any opening for chlorophyll or carotenoid pigment binding (Fan et al., 2015). These structural features align with earlier findings that PsbS, isolated from plants and reconstituted with pigments in vitro, lacks pigment-binding capacity (Dominici et al., 2002; Bonente et al., 2008). It is worth mentioning, however, that the general purification procedure cannot completely remove pigments from PsbS (Funk et al., 1995; Aspinall-O’Dea et al., 2002), and a putative chlorophyll density is observed in the crystal structure (Fan et al., 2015). Moreover, other members of the LHC superfamily, which share the LHC motif but have a much simpler structure than PsbS, such as one-helix proteins and two-helix stress-enhanced proteins, also exhibit pigment-binding capacity (Hey et al., 2017; Hey and Grimm, 2020). Thus, we cannot exclude the possibility that PsbS can interact with pigments in the thylakoid membrane.

Due to the ambiguity of its potential interaction with pigments, PsbS is currently considered to play an allosteric role in the qE mechanism, inducing conformational changes and generating quenching sites in other pigment-binding proteins, such as LHCII. Several LHCII proteins are positioned at specific locations around a dimeric PSII core complex, forming the PSII–LHCII supercomplex, a basic functional unit that regulates the photosynthetic light harvesting for water oxidation (Iwai et al., 2024). Two monomeric LHCIIs, CP26 (encoded by *LHCB5*) and CP29 (encoded by *LHCB4*), are most tightly bound to PSII core subunits (Mishra and Ghanotakis, 1994): the former with CP43 and PsbZ, and the latter with CP47 and PsbH (Boekema et al., 1995; Wei et al., 2016). In addition, CP26 interacts with a trimeric S-LHCII (S for <u>s</u>trongly bound), which also directly binds to the PSII core complex through PsbW. On the other hand, CP29 connects with another monomeric LHCII, CP24 (*LHCB6*), as well as another trimeric M-LHCII (M for <u>m</u>oderately bound) (Boekema et al., 1999a; Su et al., 2017), forming a hetero-oligomeric antenna complex, called B4C (Bassi and Dainese, 1992; Betterle et al., 2009). Interestingly, the dissociation of B4C is correlated with NPQ capacity (Betterle et al., 2009). In the dark, B4C remains associated, but HL induces its dissociation into three individual LHCII proteins. Furthermore, HL-induced B4C dissociation requires lumenal acidification, zeaxanthin accumulation, and PsbS protonation (Betterle et al., 2009). As part of PSII–LHCII supercomplexes, the dissociation of B4C in HL could indicate that NPQ induction involves the reorganization of protein–protein interactions (Johnson et al., 2011). In fact, the lack of B4C components (i.e., CP24 and CP29) reduces the level of association between PSII–LHCII supercomplexes (de Bianchi et al., 2011; Goral et al., 2012a), implying that the structure of B4C may play an important role in organizing PSII–LHCII supercomplexes into an ordered megacomplex array (Boekema et al., 1999b; Shan et al., 2024). Intriguingly, the absence of PsbS leads to a more frequent ordered array formation, whereas PsbS overexpression prevents its formation, suggesting that PsbS also influences interactions between PSII–LHCII supercomplexes (Kereïche et al., 2010; Goral et al., 2012a). Nevertheless, how PsbS protonation remodels protein–protein interactions to create the qE site remains unknown.

In this study, we revisited the *Arabidopsis* PsbS mutant, *npq4-E122Q*-*E226Q* (hereafter *E122Q-E226Q*), in which the glutamate residues at positions 122 and 226 of PsbS are substituted with glutamine (Li et al., 2004), to investigate how the physical presence of a ‘non-functional’ PsbS variant influences qE induction in relation to zeaxanthin accumulation. While consistent with previous reports that *E122Q*-*E226Q* exhibits a severe defect in NPQ induction, we show here that this mutant PsbS still allows for a small but distinctively rapid qE induction once zeaxanthin has sufficiently accumulated. Our data indicate that this small induction is strongly correlated with lumenal acidification, suggesting that the E122Q and E226Q substitutions uncover additional sites for protonation and function of PsbS in inducing qE, which becomes visible via a process directly involving zeaxanthin. Furthermore, we identify a strong correlation between this process and the presence of CP29. By using PsbS protein variants alongside null mutants that completely lack the protein, this study demonstrates how maintaining essential protein-protein interactions while eliminating specific biochemical functions can reveal molecular mechanisms that are otherwise obscured in null-mutant analyses. We will discuss how this observed small qE induction serves as a unique readout for the interaction between PsbS and zeaxanthin in vivo.

## RESULTS

### PsbS mutant affecting proton sensing can induce energy-dependent quenching in the presence of zeaxanthin

We compared the NPQ kinetics of WT, PsbS null mutant (*npq4*), and *E122Q*-*E226Q* leaf discs using two consecutive cycles of actinic light (AL) illumination and dark. The first cycle showed NPQ activation after dark acclimation, while the second cycle indicated NPQ induction following zeaxanthin accumulation (Andersson et al., 2001). Consistent with previous findings, *npq4* exhibited a severe defect in NPQ induction in both cycles compared to WT (**Fig. 1A**). Similarly, *E122Q-E226Q* showed the same NPQ defect as *npq4* in the first light-dark cycle; however, in the second cycle, we observed a rapid, although small, NPQ induction that did not occur in *npq4* (**Fig. 1A, asterisk**). A critical distinction between the two mutants is that *npq4* completely lacks PsbS proteins, whereas *E122Q-E226Q* contains the mutated PsbS variant. Therefore, despite the lack of protonation at the E122 and E226 residues, which prevents qE in the first cycle, the presence of the mutant PsbS protein and the zeaxanthin presumably accumulated during the first cycle resulted in a rapid NPQ induction in the second cycle. We termed this small but rapid NPQ response the ’*EQ* jump,’ as it was observed only in the E-to-Q amino acid substitution mutant.

**Figure 1.**
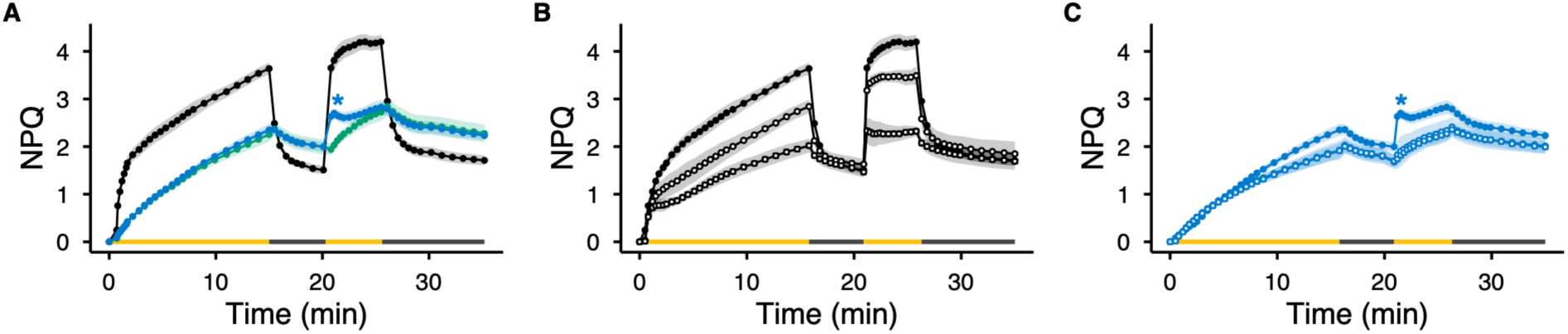
Observation of the *EQ* jump during two cycles of AL and dark in NPQ measurements. **A)** NPQ induction and relaxation kinetics over two consecutive light/dark cycles were analyzed in WT (black), *npq4* (green), and *E122Q-E226Q* (blue). **B** and **C)** NPQ kinetics were measured after 30 min of DTT treatment at 1 mM (open circle) and 5 mM (open square) concentrations for WT (B) and *E122Q-E226Q* (C), respectively. The yellow and black bars indicate the durations of the AL and dark periods. The intensity of AL was ∼1000 µmol photons m^−2^ s^−1^. An asterisk (*) indicates when the *EQ* jump occurs. All data points are reported as means ± SD (n = 9).

To investigate whether the *EQ* jump is dependent on zeaxanthin accumulation, we treated leaf discs with dithiothreitol (DTT), which inhibits ascorbate-dependent VDE activity by reducing disulfide bonds required for its function (Yamamoto and Kamite, 1972). Control measurements using WT showed that 1 mM DTT treatment significantly reduced NPQ induction, consistent with previous reports (**Fig. 1B**). With 5 mM DTT treatment, the magnitude of NPQ induction in WT was reduced to levels nearly as low as those observed in *npq4*. For *E122Q-E226Q*, 1 mM DTT treatment was sufficient to completely abolish the *EQ* jump, resulting in NPQ kinetics similar to those of *npq4* (**Fig. 1C**). This suggests that zeaxanthin accumulation is required for the *EQ* jump to occur in *E122Q-E226Q*. Although reducing agents such as DTT can interfere with other enzyme activities, it has been shown that the DTT concentration used here (1 mM) does not affect photosynthetic oxygen evolution activity (Bilger et al., 1989).

To confirm the correlation between zeaxanthin accumulation and the *EQ* jump, we determined the changes in the de-epoxidation state (DES) of xanthophylls within leaf discs using HPLC pigment analysis. The results for WT, *npq4*, and *E122Q-E226Q* indicated that the conversion from violaxanthin to zeaxanthin started within 1 min after the onset of AL illumination, and the DES for all three lines reached above 50% after 10 min of AL illumination (**Fig. 2A**). This confirms that zeaxanthin accumulation occurs similarly regardless of the presence of PsbS or the lack of WT-level NPQ induction, and that zeaxanthin had accumulated equally at the onset of the second NPQ induction. We also determined the pigment composition of the DTT-treated leaf discs (**Figs. 1B–C**). The results show that there was no significant increase in zeaxanthin accumulation due to DTT-mediated inhibition of VDE activity (**Fig. 2B**), indicating that the absence of the *EQ* jump in *E122Q-E226Q* is correlated with the lack of zeaxanthin accumulation.

**Figure 2.**
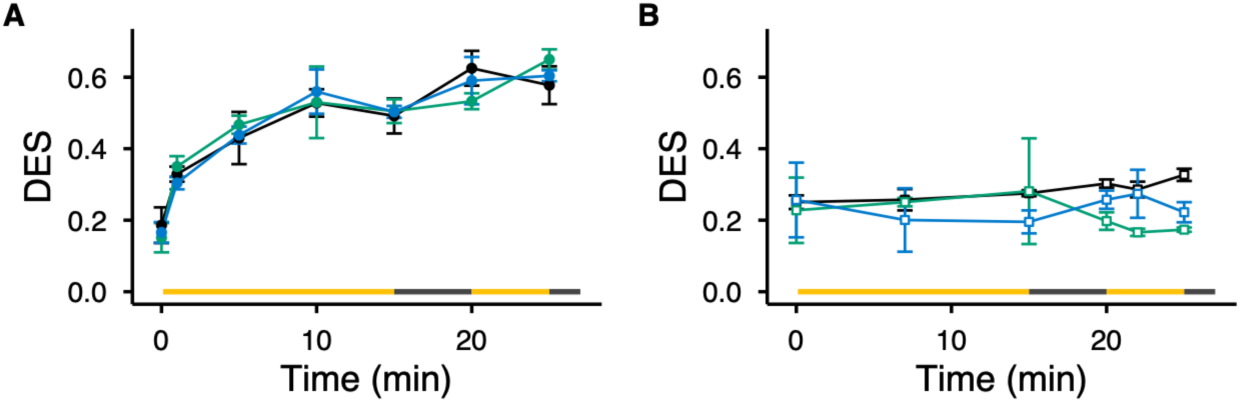
Changes in DES during NPQ measurements. **A)** DES of leaf discs was measured by HPLC pigment analysis during NPQ induction and relaxation through two consecutive cycles of AL illumination and dark in WT (black), *npq4* (green), and *E122Q-E226Q* (blue), as shown in Figure 1. **B)** The DES was measured after 30 min of DTT treatment at 5 mM concentration (open square), as shown in Figure 1. The yellow and black bars indicate the durations of the AL and dark periods. The intensity of AL was ∼1000 µmol photons m^−2^ s^−1^. All data points are reported as means ± SD (n = 3).

To further verify this correlation, we varied the duration of the first AL illumination to assess how zeaxanthin accumulation relates to the induction of the *EQ* jump. The results showed that 1 or 5 min of the first AL illumination (∼1000 µmol photons m^-2^ s^-1^) was insufficient to induce the *EQ* jump in *E122Q-E226Q*, whereas rapid NPQ induction and relaxation were observed in WT, reconfirming the strong contribution of PsbS protonation to rapid NPQ kinetics (**Figs. 3A, B**). To induce a noticeable *EQ* jump in *E122Q-E226Q*, 10 min of AL illumination was required to allow sufficient zeaxanthin accumulation (**Fig. 3C, arrow**). Based on our pigment analysis (**Fig. 2A**), 1 min and 5 min of AL illumination generated a DES of approximately 30% and 40%, respectively, whereas a DES above 50% was observed after 10 min of AL illumination. This indicates that at least half of the violaxanthin in the thylakoid needs to be converted to antheraxanthin/zeaxanthin to trigger the *EQ* jump in *E122Q-E226Q*. Because the absence of PsbS in *npq4* precluded the *EQ* jump, our findings suggest that the presence of the *E122Q-E226Q* PsbS variant in the thylakoid membrane is necessary for the *EQ* jump to occur once zeaxanthin has sufficiently accumulated.

**Figure 3.**
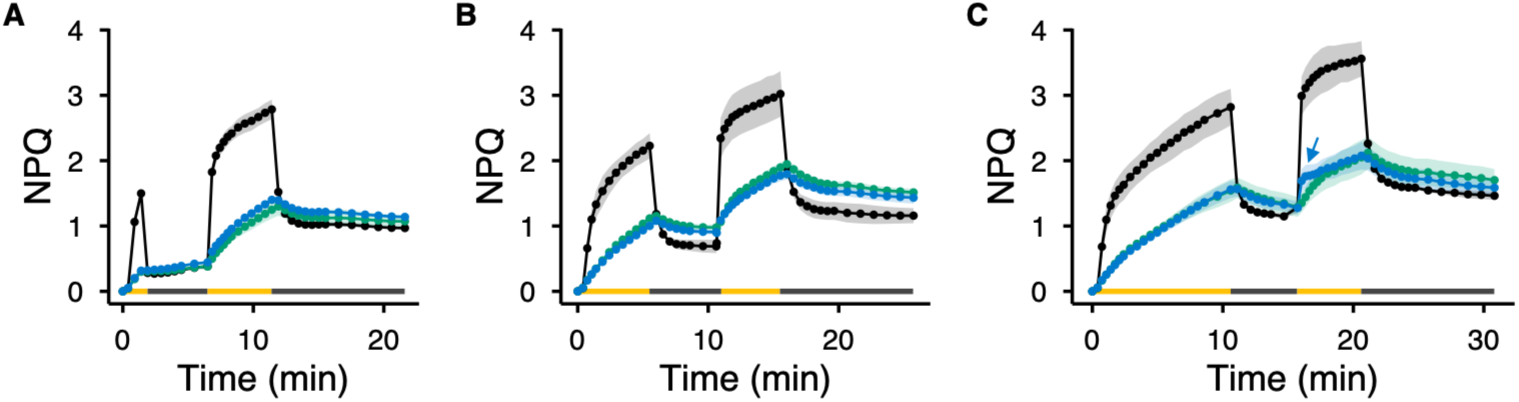
Effect of the initial AL duration on *EQ* jump induction during two light/dark cycles in NPQ measurements. NPQ induction and relaxation kinetics were analyzed in WT (black), *npq4* (green), and *E122Q-E226Q* (blue). The duration of the first AL was varied: **A)** 1 min, **B)** 5 min, and **C)** 10 min. The yellow and black bars indicate the durations of the AL and dark periods. The intensity of AL was ∼1000 µmol photons m^−2^ s^−1^. An arrow indicates when the *EQ* jump became visible. All data points are reported as means ± SD (n = 3).

Next, to examine whether the *EQ* jump is caused by light-induced lumenal acidification, we observed NPQ kinetics after nigericin treatment, which dissipates the proton gradient across the thylakoid membrane (ΔpH) by exchanging H^+^ and K^+^ (Wraight and Crofts, 1970). The results indicated that NPQ induction was severely hindered for both WT and *E122Q-E226Q*, and the absence of lumenal acidification abolished the *EQ* jump in the latter (**Fig. 4A–B**). However, nigericin treatment also inhibits VDE, which is also activated by protonation (Bugos and Yamamoto, 1996; Arnoux et al., 2009). To determine the specific contribution of light-induced lumenal acidification to the *EQ* jump, we lowered the AL intensity on the second NPQ induction cycle, thereby developing lower levels of lumenal acidification. The results demonstrate that the extent of the *EQ* jump became smaller when lower AL intensities were used, even though zeaxanthin was sufficiently accumulated after the first cycle (**Fig. 4C**). Therefore, the induction of the *EQ* jump is caused by lumenal acidification, suggesting that protonation exists in PsbS at sites other than E122 and E226.

**Figure 4.**
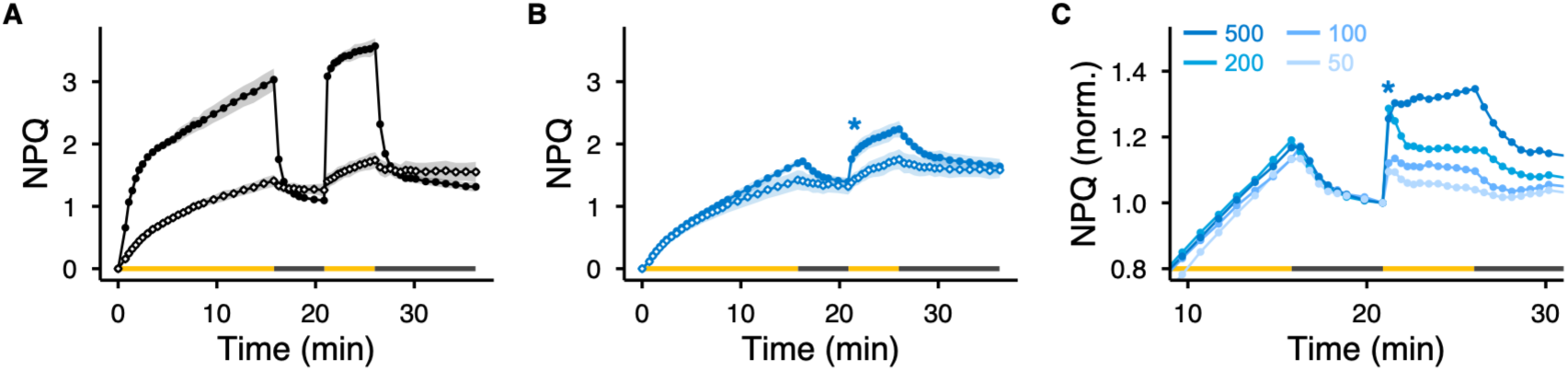
Correlation between light-induced lumenal acidification and *EQ* jump induction during two light/dark cycles in NPQ measurements. NPQ induction and relaxation kinetics over two consecutive light/dark cycles were analyzed for control measurements (closed circle) in **A)** WT and B) *E122Q-E226Q*. NPQ kinetics were measured after 30 min of nigericin treatment at 50 µM (open diamond). The intensity of AL was ∼1000 µmol photons m^−2^ s^−1^. **C)** NPQ kinetics were analyzed in *E122Q-E226Q,* with varying second AL intensity: 50, 100, 200, and 500 µmol photons m^−2^ s^−1^ (indicated by numbers). The first AL intensity was ∼1000 µmol photons m^−2^ s^−1^. The graph shows a focused view of the *EQ* jump, with SD shading omitted for clarity. For direct comparison, the data were normalized at the end of the first dark period. An asterisk (*) indicates when the *EQ* jump occurs. The yellow and black bars indicate the durations of the AL and dark periods. All data points are reported as means ± SD (n = 3).

The *EQ* jump reflects an additional increase in NPQ at the onset of the second AL illumination in *E122Q-E226Q*, a phenomenon that is absent in *npq4*. We thus examined whether extending the duration of the first AL, without a light/dark cycle, would increase NPQ induction in *E122Q-E226Q* relative to *npq4*. Interestingly, extended AL exposure resulted in very similar NPQ kinetics in both mutants (**Fig. 5A**). The only difference in this experimental condition was the absence of a dark-to-light shift, as prolonged AL exposure allows for sufficient zeaxanthin accumulation along with the development of a ΔpH. This suggests that the *EQ* jump becomes apparent only upon a sudden change in light-induced lumenal acidification. Next, we examined whether the magnitude of the *EQ* jump increases with additional light/dark cycles. While the total NPQ caused by the *EQ* jump increased with each cycle, the steady-state NPQ levels in both *E122Q-E226Q* and *npq4* were the same after 3 min of each AL illumination (**Fig. 5B**). These observations reconfirm the essential role of the two glutamate residues (E122 and E226) of PsbS in achieving high NPQ induction. Moreover, the *EQ* jump, only visible in the presence of the mutant PsbS protein, reveals that the *E122Q-E226Q* mutant is still weakly functional in the presence of zeaxanthin.

**Figure 5.**
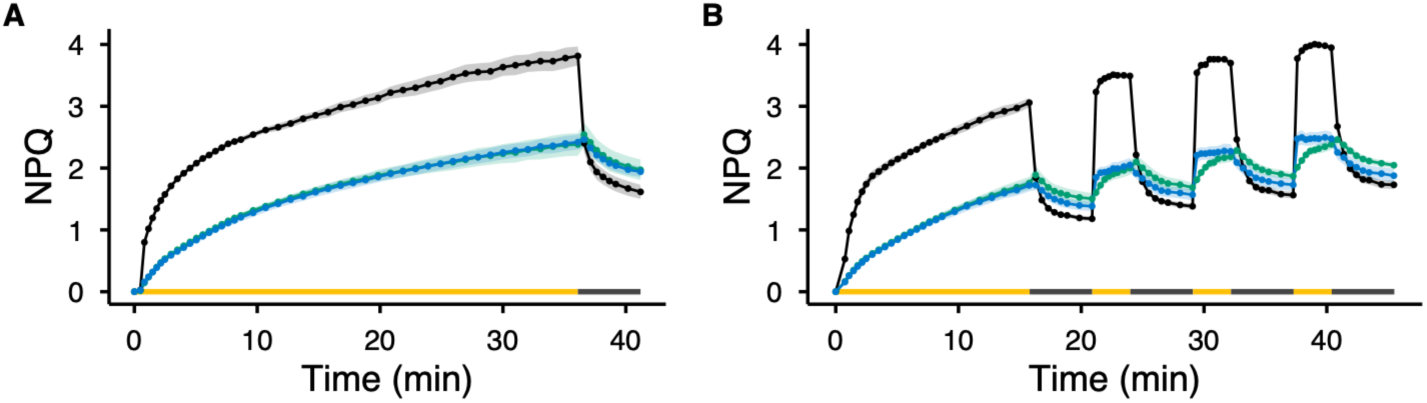
NPQ induction kinetics under altered light/dark cycles. NPQ induction and relaxation kinetics were analyzed in WT (black), *npq4* (green), and *E122Q-E226Q* (blue). **A)** The duration of the first AL was extended to 35 min, followed by 5 min of dark. **B)** After the initial light/dark cycle, 3 min of AL and 5 min of dark were repeated three times. The yellow and black bars indicate the durations of the AL and dark periods. The intensity of AL was ∼1000 µmol photons m^−2^ s^−1^. All data points are reported as means ± SD (n = 3).

### PsbS protonation at E122 and E226 differentially affects the *EQ* jump kinetics

Based on our observations, we identified three factors for the *EQ* jump to occur: the presence of the mutant PsbS protein (**Fig. 1**), zeaxanthin accumulation (**Figs. 1–3**), and light-induced lumenal acidification (**Figs. 4–5**). Next, we investigated how each of the glutamates (E122 and E226) influences the *EQ* jump by using the two single-protonation mutants, *npq4-E122Q* and *npq4-E226Q* (hereafter *E122Q* and *E226Q*, respectively) (Li et al., 2004). Consistent with previous literature, both *E122Q* and *E226Q* exhibited higher levels of NPQ induction than *E122Q-E226Q* but lower levels than WT during the first AL period (**Fig. 6A**). At the onset of the second AL illumination, however, each mutant showed steep NPQ induction, indicating that each of the two glutamate residues contributes significantly to the rapid response to the dark-to-light shift (**Fig. 6A**). Intriguingly, the NPQ kinetics during the second AL period were quite distinct between the two mutants: *E122Q* showed a small dip in NPQ and remained similar for the rest of the AL period, whereas *E226Q* showed a steep rise and fall of NPQ. These rapid qE kinetics for both *E122Q* and *E226Q* remained sensitive to DTT treatment, although a higher concentration was required to diminish the qE compared with *E122Q-E226Q* (**Figs. 6B–C**). HPLC pigment analysis showed that zeaxanthin accumulation occurred in both mutants, similar to that observed in *E122Q-E226Q* (**Supplementary Fig. S1**). Also, varying the duration of the first AL period influenced the magnitude of rapid qE induction at the onset of the second AL illumination (**Supplementary Fig. S2**), indicating that light-induced zeaxanthin accumulation is involved in this process. Moreover, the absence of lumenal acidification by nigericin treatment also abolished this process (**Supplementary Fig. S3**). Therefore, while the rapid qE induction caused by single protonation at E122 or E226 shares the same mechanisms as the *EQ* jump, the mutants affecting each of the two protonatable residues exhibited distinct NPQ kinetics during the second AL period. The crystal structure of purified spinach PsbS forms a dimer (Fan et al., 2015), in which E122 (equivalent to E69 of spinach PsbS) is facing outward, while E226 (equivalent to E173 of spinach PsbS) is facing the interface of the dimer. The transition between monomeric and dimeric forms of PsbS upon lumenal acidification and their potential role in NPQ have been discussed in the literature (for example, see discussion in van Amerongen and Croce, 2025). Despite the uncertainty of the PsbS monomer-dimer model, the distinct kinetics of the zeaxanthin-dependent qE induction between *E122Q* and *E226Q* suggest different functional roles for each protonation site in NPQ induction.

**Figure 6.**
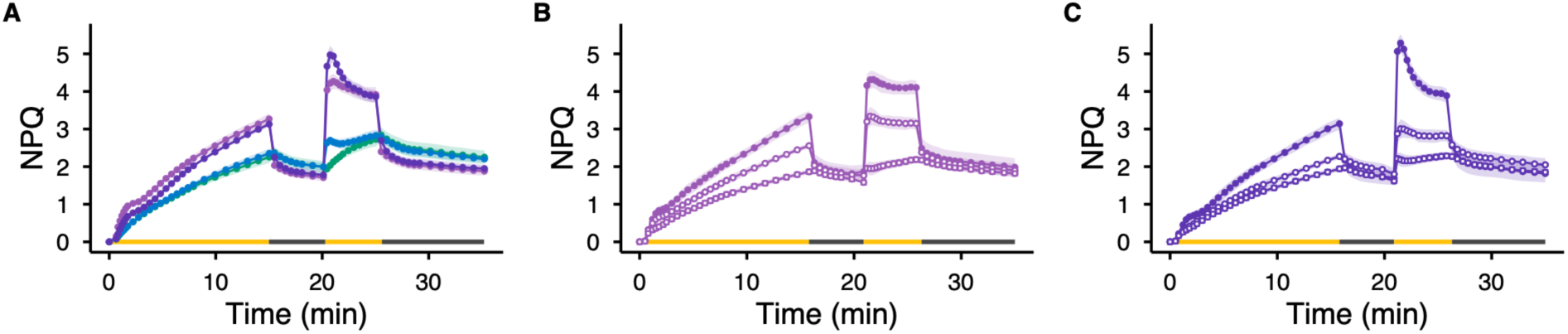
The effect of the single-protonation mutation in PsbS on the *EQ* jump during two cycles of AL and dark in NPQ measurements. **A)** NPQ induction and relaxation kinetics over two consecutive light/dark cycles were analyzed in *npq4* (green), *E122Q-E226Q* (blue), *E122Q* (light purple), and *E226Q* (dark purple). **B** and **C)** NPQ kinetics were measured after 30 min of DTT treatment at 1 mM (open circle) and 5 mM (open square) concentrations for *E122Q* (B) and *E226Q* (C), respectively. The yellow and black bars indicate the durations of the AL and dark periods. The intensity of AL was ∼1000 µmol photons m^−2^ s^−1^. All data points are reported as means ± SD (n = 9).

### The absence of B4C formation reduces the magnitude of the *EQ* jump

The formation of B4C, a hetero-oligomeric antenna complex composed of CP29, CP24, and M-LHCII, has been shown to remain stable in the dark, but it dissociates upon NPQ induction (Betterle et al., 2009). B4C disassembly is correlated with HL, lumenal acidification, and zeaxanthin accumulation. Interestingly, in the two PsbS single-protonation mutants, *E122Q* and *E226Q*, B4C components were partially dissociated, whereas no disassembly occurred in the double-protonation mutant, *E122Q-E226Q* (Betterle et al., 2009), suggesting that PsbS protonation is necessary for B4C disassembly. To investigate how the absence of B4C formation affects the *EQ* jump, we generated the *E122Q-E226Q koLhcb4* mutant, which contains the PsbS variant but lacks CP29. Because the absence of CP29 destabilizes CP24 (de Bianchi et al., 2011), B4C formation does not occur in this mutant. NPQ measurements showed that the absence of CP29 significantly reduced both the overall magnitude of NPQ and the extent of the *EQ* jump compared to *E122Q-E226Q* (**Fig. 7A**). HPLC pigment analysis revealed that zeaxanthin accumulation in the *E122Q-E226Q koLhcb4* mutant was similar to that observed in *E122Q-E226Q* (**Supplementary Fig. S4**). To more clearly observe the maximum potential of the *EQ* jump, the first AL period was extended to 30 min to increase zeaxanthin accumulation (**Fig. 7B**). Using this extended illumination, we compared the *EQ* jump across different lines by normalizing the magnitude to a relative scale (**Table 1**). With functional PsbS present, WT and *koLhcb4* showed initial NPQ values of 0.52 ± 0.06 and 0.58 ± 0.08, respectively, at the beginning of the second AL period, whereas *npq4* exhibited a value of 0.06 ± 0.01 due to the absence of PsbS (**Fig. 7B**, **Table 1**). On the other hand, after 30 min of AL illumination and higher zeaxanthin accumulation, the *EQ* jump observed in the *E122Q-E226Q* reached 0.31 ± 0.01, while the absence of CP29 in the *E122Q-E226Q koLhcb4* mutant caused a decrease to 0.15 ± 0.04 (**Table 1**). This demonstrates that the absence of CP29 negatively impacts the magnitude of the *EQ* jump. We also compared how the magnitude of the *EQ* jump changed during repeated light/dark cycles, as shown in **Fig. 5B**. In the presence of functional PsbS, WT and *koLhcb4* reached their highest NPQ values first (at 21 min) and maintained similar levels throughout the cycles, whereas *npq4* remained low due to the lack of PsbS (**Figs. 5B and 7B**, **Table 2**). In contrast, the magnitude of the *EQ* jump in *E122Q-E226Q* increased slightly during the repeated cycles, but remained low in the *E122Q-E226Q koLhcb4* mutant (**Fig. 7B**, **Table 2**). This observation confirms that the presence of CP29 significantly contributes to the magnitude of the *EQ* jump.

**Figure 7.**
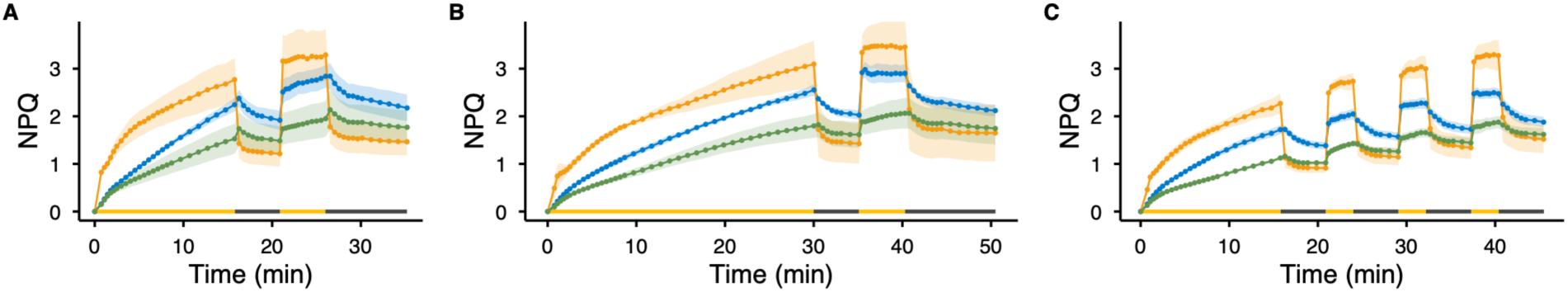
The effect of the absence of CP29 on the *EQ* jump. NPQ induction and relaxation kinetics were analyzed in *E122Q-E226Q* (blue), *koLhcb4* (orange), and *E122Q-E226Q koLhcb4* (sage green). **A)** NPQ induction and relaxation kinetics over two consecutive light/dark cycles were analyzed. **B)** The duration of the first AL was extended to 30 min, followed by 5 min of dark, 5 min of the second AL, and 10 min of dark. **C)** After the initial light/dark cycle, 3 min of AL and 5 min of dark were repeated three times. The yellow and black bars indicate the durations of the AL and dark periods. The intensity of AL was ∼1000 µmol photons m^−2^ s^−1^. All data points are reported as means ± SD (n = 3).

**Table 1.** The magnitude of rapid qE induction after 30 min of AL illumination (Fig. 7B).

|  | WT | *npq4* | *E122Q-E226Q* | *koLhcb4* | *E122Q-E226Q*<br>*koLhcb4* |
|---|---|---|---|---|---|
| at 35 min | 0.52 ± 0.06 | 0.06 ± 0.01 | 0.31 ± 0.01 | 0.58 ± 0.08 | 0.15 ± 0.04 |

The magnitude of rapid qE induction was calculated as the difference in NPQ values (before and 20 sec after the onset of the second AL illumination) divided by the NPQ value taken after the illumination. The values are reported as means ± SD (n = 3).

**Table 2.** The magnitude of rapid qE induction during three consecutive light/dark cycles (Figs. 5B & 7C).

|  | WT | *npq4* | *E122Q-E226Q* | *koLhcb4* | *E122Q-E226Q koLhcb4* |
|---|---|---|---|---|---|
| at 21 min | 0.63 ± 0.02 | 0.06 ± 0.02 | 0.25 ± 0.02 | 0.63 ± 0.03 | 0.16 ± 0.01 |
| at 29 min | 0.61 ± 0.02 | 0.07 ± 0.01 | 0.29 ± 0.01 | 0.60 ± 0.04 | 0.18 ± 0.02 |
| at 37 min | 0.59 ± 0.02 | 0.07 ± 0.01 | 0.30 ± 0.02 | 0.58 ± 0.05 | 0.18 ± 0.02 |

The magnitude of rapid qE induction was calculated as the difference in NPQ values (before and 20 sec after the onset of each AL illumination) divided by the NPQ value taken after each of the AL illuminations. The values are reported as means ± SD (n = 3).

## DISCUSSION

### Additional protonation sites other than E122 and E226 exist in PsbS

Previous experiments identified the two glutamate residues, E122 and E226, as essential protonation sites for *Arabidopsis* PsbS to induce qE (Li et al., 2004). Sequence-wise, these two residues are symmetrically positioned within the TM1–2 and the TM3–4 loops. However, crystal structure and AlphaFold predictions suggest that these lumenal loop domains are not structurally symmetrical (Fan et al., 2015; see PDB ID 4RI2 and AF ID Q9XF91 for details). Specifically, the TM1–2 loop contains two short amphipathic helices, called H1 and H3, which are four and three amino acids long, respectively, while the TM3–4 loop contains a long amphipathic helix, called H2, which is nine amino acids long (Fan et al., 2015; Liguori et al., 2019). E122 is located between H1 and H3, and E226 is located within H2. Molecular dynamics simulations showed that while the H1 helix transitioned between a 3_10_-helix and a turn with no observable pH dependence, the H3 helix was highly pH-sensitive, completely unfolding at high pH (7 and 8) and shifting from a 3_10_- helix to a turn/coil (Liguori et al., 2019). Coarse-grained simulations revealed that the H2 helix also undergoes a repositioning from the aqueous to the membrane phase upon a transition from neutral to acidic pH (Chiariello et al., 2023), experimentally supported by vibrational spectroscopy (Krishnan-Schmieden et al., 2021). Therefore, the light-induced protonation at both E122 and E226 triggers a structural shift across all three lumenal amphipathic helices. Importantly, the substitution of these two residues abolishes qE, indicating that PsbS protonation is a critical factor for the rapid response to changing light conditions (Li et al., 2004).

In this study, we revisited the NPQ kinetics of the *Arabidopsis* mutant line *E122Q-E226Q*, which contains PsbS with amino acid substitutions (E122Q and E226Q) in the *npq4* background (Li et al., 2004). Previous reports indicated that initial NPQ induction upon AL illumination is similar to that of *npq4* (a null PsbS mutant), suggesting a deficiency in qE due to the lack of protonation at E122 and E226 (Li et al., 2000; Li et al., 2004). However, we found that *E122Q-E226Q* is not completely non-functional, as it still induces a rapid but small qE upon dark-to-light shifts after zeaxanthin has accumulated, a phenomenon we term the *EQ* jump (**Figs. 1–5**). The small magnitude of the *EQ* jump, compared to a regular qE induction, is primarily due to the absence of fully functional PsbS. In fact, the single-protonation mutants, *E122Q* and *E226Q*, showed significant restoration of the magnitude of NPQ, especially for the zeaxanthin-dependent rapid qE induction with distinct kinetics (**Fig. 6**). These observations reconfirm that the rapid response to changing light conditions depends significantly on the presence of PsbS and its protonation upon lumenal acidification.

The observation of the *EQ* jump implies the potential existence of other protonation sites within PsbS, whose role becomes more evident following zeaxanthin accumulation. Aside from E122 and E226, PsbS possesses six other protonatable residues on the lumenal side: E108, E129, E131, E212, D233, and E235. Similar to E122 and E226, these residues are pseudo-symmetrically distributed within the TM1–2 and TM3–4 loops, pairing as follows: E108 with E212, E129 with D233, and E131 with E235. Earlier site-directed mutagenesis studies have shown that, for the symmetrical pair E108 and E212, mutating a single residue to glutamine did not affect PsbS protein accumulation, but mutating both residues abolished it, likely due to a loss of protein stability, as both residues are located at the end of the α-helices of TM1 and TM3 (Li et al., 2002b). For the pair E129 and D233, double mutation did not alter qE levels compared to the control. However, substituting E129 with glutamine caused a slight increase in qE, while the D233N mutation resulted in a ∼50% reduction in qE (Li et al., 2002b). These differences may be due to the structurally asymmetric relationship between the two, with E129 located within the H3 helix and D233 situated at the edge of the TM4 helix on the lumenal side. Similarly, single-residue substitutions of E131 and E235 to glutamine caused ∼34% and ∼10% reductions in qE, respectively, which were not attributed to changes in protein stability or expression levels. E131 is located near the H3 helix, while E235 is at the TM4 helix, and mutating both residues caused a ∼50% reduction in qE (Li et al., 2002b). These results imply complex functional roles of each possible protonation site, as E129 and E131 (near the H3 helix) appear to impact qE positively and negatively, respectively. Also, while both D233 and E235 are situated at the TM4 helix, D233N causes a 50% reduction in qE, whereas E235Q appears less affected. Ultimately, the definitive identification of the sites responsible for additional protonation will require thorough analysis of varied combinations of substitution mutations in future work.

Intriguingly, observing the *EQ* jump requires a transient high ΔpH in the presence of zeaxanthin, and it is not observed during extended AL illumination, despite the presence of a stable ΔpH and the accumulation of what would otherwise be sufficient zeaxanthin (**Fig. 5A**). When NPQ kinetics under continuous AL (**Fig. 5A)** are compared with those under repeated light/dark cycles (**Fig. 5B**), the *EQ* jump rapidly reaches the NPQ level observed under continuous light conditions (see arrows in **Supplementary Fig. S5** for direct comparison). This indicates that PsbS maintains a conformation and/or location that allows for a rapid response to sudden HL exposure. Molecular dynamics simulations previously calculated the pKa values for each protonatable residue in PsbS. The two essential protonatable residues, E122 and E226 (equivalent to E69 and E173 in spinach PsbS), have pKa values of 5.2 and 6.1, respectively (Liguori et al., 2019). Notably, E212 (E159 in spinach) has the lowest pKa value of 4.1 among the residues on the lumenal side (Liguori et al., 2019). As mentioned above, the single E212Q mutation does not affect qE, presumably because E122/E226 protonation remains the primary trigger, whereas in the *E122Q-E226Q* background, E212 protonation could contribute to the *EQ* jump.

Several other residues exhibit slightly higher pKa values: E129 (E76 in spinach) at 5.7, E131 (E78 in spinach) at 6.3, and D233 (E180 in spinach), whose pKa remains undetermined in *Arabidopsis* but is presumably comparable to the spinach pKa value of 6.4 (Liguori et al., 2019). As discussed above, earlier substitution experiments suggest that these additional protonatable residues may contribute to qE in ways distinct from the E122/E226 trigger. Importantly, these residues with slightly higher pKa values show substantial pKa shifts relative to free glutamate in water (pKa = 4.0–4.3; Forsyth et al., 2002; Grimsley et al., 2009), indicating that they remain protonatable within the range of lumenal pH shifts. Because their pKa values are higher than those of E122 and E226, these residues would remain protonated longer when lumen pH rises during the dark period. Additionally, given their proximity to the TM helices, this prolonged protonation may stabilize lumenal domains in a standby conformation without triggering quenching, consistent with the observation that *E122Q-E226Q* does not show higher NPQ relative to *npq4* under continuous AL (**Fig. 5A**). At the onset of sudden HL exposure, however, the combination of zeaxanthin accumulation and a transiently high ΔpH may rapidly activate this standby state, producing the *EQ* jump, which reaches the NPQ level observed under continuous AL but cannot exceed it without the E122/E226 trigger (**Supplementary Fig. S5**). Consistent with this hypothesis, coarse-grained simulations appear to support a prolonged protonated state for E131 in particular (Chiariello et al., 2023).

### The *EQ* jump as a readout for interplay between PsbS, zeaxanthin, and CP29

Based on our observations, visualizing the *EQ* jump requires three factors: the presence of PsbS protein, zeaxanthin accumulation, and light-induced lumenal acidification (**Fig. 8**). If PsbS is absent from the thylakoid membrane (e.g., *npq4*), qE does not occur (**Fig. 1A**). Similarly, if PsbS is present but the proton gradient from light-induced lumenal acidification is not established (e.g., by nigericin treatment), qE does not occur (**Fig. 4A**). When PsbS senses the proton gradient but zeaxanthin accumulation is insufficient (e.g., by DTT treatment), the magnitude of qE is significantly reduced compared to normal qE (**Fig. 1B**). On the other hand, when PsbS lacks full proton-sensing capacity (e.g., *E122Q-E226Q*) but zeaxanthin is sufficiently accumulated, the *EQ* jump is observed (**Figs. 1–2**). Importantly, the magnitude of the *EQ* jump is strongly correlated with both zeaxanthin accumulation and lumenal acidification levels (**Figs. 3–4**).

**Figure 8.**
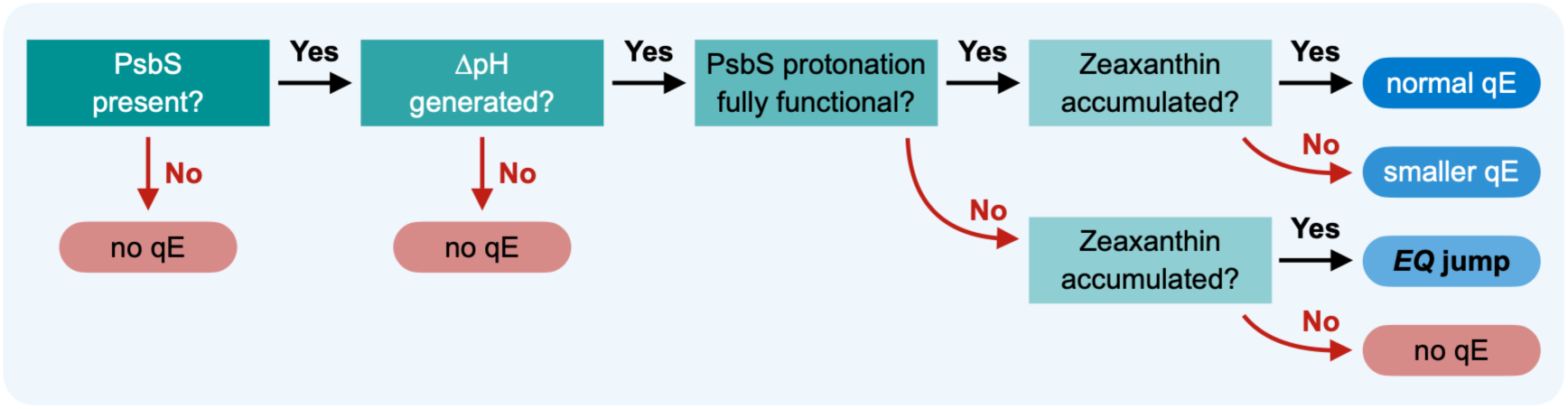
Conditions required for the *EQ* jump. For this figure, qE is defined specifically as the rapid NPQ induction induced immediately upon a dark-to-light transition. All other factors, including pigment and protein compositions, are assumed equal.

Not only does the mechanism by which PsbS protonation causes qE remain unclear, but it has also long been debated whether zeaxanthin contributes to qE directly or allosterically. Several lines of evidence suggest a specific role of zeaxanthin in qE: i) a strong correlation exists among zeaxanthin accumulation, lumenal acidification, and qE induction (Gilmore and Yamamoto, 1993); ii) the absence of zeaxanthin, induced by DTT treatments or by genetic mutations in VDE, hinders qE induction (Yamamoto and Kamite, 1972; Bilger et al., 1989; Niyogi et al., 1998); iii) the formation of a zeaxanthin S_1_ state for excitation energy transfer (EET) quenching or zeaxanthin radical cation for charge transfer (CT) quenching is observed in the thylakoid membrane, which is strongly correlated with qE induction (Holt et al., 2005; Lee et al., 2024); and iv) CT quenching is observed in CP29, CP26, and CP24, correlated with the presence of zeaxanthin (Ahn et al., 2008; Avenson et al., 2008).

As another xanthophyll bound to LHCII, lutein has also been considered a direct quencher involved in qE (Ruban, 2016; van Amerongen and Croce, 2025). Unlike zeaxanthin, which is generated through the reversible conversion of violaxanthin by light-induced VDE activity, lutein is constitutively present in the thylakoid membrane and situated within the core of the LHCII protein at the binding sites called L1 and L2 (Liu et al., 2004; Pan et al., 2011). To prevent constitutive dissipation of absorbed light energy, lutein would need to switch between light-harvesting and quenching states. Several mechanisms have been proposed for such a switch: i) EET from chlorophylls to the S_1_ state of lutein (Ruban et al., 2007; Chmeliov et al., 2015; Balevičius et al., 2017); ii) excitonic interactions between chlorophylls and the S_1_ state of lutein (van Amerongen and van Grondelle, 2001; Holleboom and Walla, 2014); and iii) EET from chlorophylls to the S* state of lutein (Liguori et al., 2017; Mascoli et al., 2019).

Further genetic studies have indicated the sites of quenching induced by zeaxanthin and lutein. *Arabidopsis NoM* mutants, which lack monomeric LHCIIs (CP24, CP26, and CP29) but retain LHCII trimers, display three distinct NPQ kinetic characterizations: (i) PsbS-dependent rapid qE induction upon AL illumination, (ii) an absence of the early phase of NPQ induction, and (iii) the presence of the later phase of NPQ induction (Dall’Osto et al., 2017). This biphasic response after AL illumination suggests that monomeric LHCIIs drive the early phase, whereas LHCII trimers are responsible for the later phase. Notably, the loss of lutein in *NoM lut2* plants led to only a marginal decrease in NPQ amplitude relative to *NoM* (Dall’Osto et al., 2017). Conversely, the lack of zeaxanthin in *NoM npq1* plants resulted in a severe impairment of NPQ induction, indicating that zeaxanthin accumulation is critical for this process within LHCII trimers (Dall’Osto et al., 2017). Moreover, studies using *Arabidopsis koLHCII* mutants, which lack LHCII trimers but retain monomeric LHCIIs, indicate that the majority of NPQ occurs in the LHCII trimers (Nicol et al., 2019; Guardini et al., 2025; Nicol et al., 2026). Surprisingly, the lack of lutein in monomeric LHCIIs in *koLHCII lut2* caused only a small decrease in NPQ amplitude relative to *koLHCII* (Guardini et al., 2025).

Notably, PsbS-dependent rapid qE induction upon AL illumination is still observed in mutants lacking either monomeric or trimeric LHCIIs (Dall’Osto et al., 2017; Nicol et al., 2019; Guardini et al., 2025). Extensive studies on the role of CP29 in NPQ have suggested that zeaxanthin binding at the L2/V1 sites or the periphery triggers formation of the quenching site (Caffarri et al., 2001; Morosinotto et al., 2002; Ahn et al., 2008; Avenson et al., 2008; Betterle et al., 2009; de Bianchi et al., 2011; Guardini et al., 2020; Guardini et al., 2025). We observed that the amplitude of the *EQ* jump decreased by 40–50% in the absence of CP29 in the *E122Q-E226Q koLhcb4* mutant (**Fig. 7**, **Tables 1–2**). Because the absence of CP29 not only destabilizes CP24 (de Bianchi et al., 2011) but also alters the macromolecular organization of PSII–LHCII supercomplexes (de Bianchi et al., 2011; Goral et al., 2012b), this reduced magnitude cannot be attributed solely to the loss of CP29. Importantly, however, the 40–50% reduction is specific to the loss of CP29 in the *E122Q-E226Q* background, since the rapid qE induction observed in *koLhcb4* is comparable to that in WT (**Tables 1–2**). This result points to a functional interaction between PsbS and CP29 (or B4C) that underlies the *EQ* jump, which would be obscured when PsbS is fully functional. Neither our data nor the available literature can confirm or rule out the allosteric effector hypothesis, since atomic-level evidence for interactions among zeaxanthin, CP29, and PsbS remains lacking. Our results nevertheless suggest that the *EQ* jump serves as a readout of PsbS–zeaxanthin interactions, potentially mediated by CP29 or B4C.

As recently noted (Nicol et al., 2026), it remains largely unclear how qE is induced following HL-induced lumenal acidification, subsequent PsbS protonation, and zeaxanthin accumulation. Whether PsbS is essential for this process remains controversial, based on results showing WT levels of NPQ in isolated chloroplasts from *npq4* when treated with an artificial mediator of cyclic electron flow (e.g., diaminodurene) to induce an unnaturally low lumen pH (Johnson and Ruban, 2011). However, it has also been demonstrated that structural changes in LHCII induced by low pH in vitro did not trigger a quenching state (Do et al., 2026), although this result may not rule out a role for specific lumenal-side protonation of LHCII under physiological lumenal acidification. Furthermore, the removal of protonation sites in CP29 did not significantly alter NPQ, suggesting that PsbS may be the sole thylakoid membrane protein that senses lumenal acidification to trigger qE induction (Guardini et al., 2020). Our observations of the *EQ* jump suggest that additional protonation sites exist within PsbS, which become evident once zeaxanthin has sufficiently accumulated. Our data from single-protonation mutants further imply that the structurally asymmetrical pairs of PsbS protonation sites play distinct roles in NPQ induction, potentially relating to a transition between dimeric and monomeric conformations. Determining the precise location of PsbS around PSII–LHCII supercomplexes will be critical to clarifying these elusive mechanisms. Recent in situ structures of PSII–LHCII supercomplexes have yet to resolve the position of PsbS (Li et al., 2026), which may be due to its low substoichiometric abundance, stochastic association with the supercomplexes, or its position distal to the supercomplex periphery.

## MATERIALS AND METHODS

### Plant strains and growth conditions

We used the following *Arabidopsis thaliana* lines for this study: Col-0 (WT), *npq4*, *npq4-E122Q-E226Q*, *npq4-E122Q*, *npq4-E226Q*, and *koLhcb4* (as described in Li et al., 2000; Li et al., 2004; de Bianchi et al., 2011). Seeds were sterilized with a bleach solution [50% (v/v) bleach, 0.025% (w/v) Tween-20] and plated on 0.5x Murashige & Skoog medium according to Arabidopsis Biological Resource Center procedures. Seeds were vernalized at 4 °C in the dark for 3 days, then germinated under a 14 h light (100 µmol photons m^-2^ s^-1^) and 10 h dark cycle with fluorescent bulbs, ambient CO_2_ (22 °C during the day and 21 °C at night). Two-week-old seedlings were transferred to Lambert LM-6 High Porosity Mix soil and grown under a 10 h light (100 µmol photons m^-2^ s^-1^) and 14 h dark cycle with fluorescent bulbs, ambient CO_2_ (23 °C during the day and 21 °C at night). Plants were bottom-watered weekly, with Miracle-Gro fertilizer applied every other watering. Experiments were conducted on 6-week-old plants using the youngest fully matured leaves.

We generated the *npq4-E122Q-E226Q koLhcb4* mutant as follows. Pollen from *koLhcb4* anthers was used to fertilize *npq4-E122Q-E226Q* carpels to generate F1 seeds. F1 plants were allowed to self-fertilize, and F2 plants were subjected to screening. We performed chlorophyll fluorescence measurements to confirm the *npq4-E122Q-E226Q* phenotype by screening for the absence of qE kinetics. The knockout of three genes encoding CP29 (*LHCB4.1*, *LHCB4.2*, and *LHCB4.3*) was confirmed by PCR genotyping. Primers used for genotyping *LHCB4* genes were previously described (de Bianchi et al., 2011) (see also **Supplementary Table S1**). Protein abundance of PsbS-E122Q-E226Q was confirmed by SDS-PAGE and immunoblot analysis using isolated thylakoid membranes with an anti-PsbS antibody (Li et al., 2000).

### Chlorophyll fluorescence measurements

To compare NPQ kinetics in multiple different lines simultaneously, we measured leaf discs floated on 300 µL of water in 96-well plates using an IMAGING-PAM (Walz). Leaf discs were dark-acclimated for 30 min before measurements. The intensity and duration of actinic light are specified in the respective figure captions. NPQ and Fv/Fm values are calculated using the following equations: NPQ = (Fm – Fm’)/Fm’; and Fv/Fm = (Fm – Fo)/Fm. DTT treatment was done during the 30-min dark acclimation by floating the leaf discs on 300 µL of 1 or 5 mM DTT in water. Nigericin treatment was done by vacuum infiltration as follows. Three leaf discs were floated in 2 mL of 50 µM nigericin in water (prepared from a 1 mM stock in 100% ethanol) in a 24-well plate and subjected to a vacuum of 400 mm Hg (17 inches) for 1 min, followed by a quick release. This process was repeated if the nigericin solution did not visibly penetrate the leaf discs.

### HPLC pigment analysis

Leaf discs were exposed to various light treatments, with and without DTT or nigericin, using the IMAGING-PAM as described above, and then flash-frozen in liquid nitrogen. Pigment extraction was performed in dim light conditions and on ice as follows. Each flash-frozen leaf disc was homogenized with Lysing Matrix D beads in 100 µL of 100% ethanol using a bead-beating grinder and lysis system (FastPrep-24 5G, MP Biomedicals) for 40 sec at 6.0 m/s, then centrifuged at 20,800 × *g* for 10 min at 4 °C. The supernatant was filtered using a 0.2 µm filter spin column at 2,700 × *g* for 3 min at 4 °C. The sample was analyzed by HPLC using a C30 reverse-phase column (YMC Carotenoid, 250 × 4.6 mm I.D., S-5 µm) with the solvent A [98% (v/v) methanol/water], solvent B [95% (v/v) methanol/water], and solvent C [100% (v/v) methyl tert-butyl ether]. The gradient was as follows (adapted from Gupta et al., 2015): a linear gradient from 100% A to 60% A and 40% C for 20 min, followed by a shift to 60% B and 40% C and then to 100% C for 2 min, and lastly re-equilibrating to 100% A for 8 min. The flow rate was 1 mL/min, and the column temperature was maintained at 30 °C. Pigment concentrations were quantified based on the peak area of each pigment detected at 445 nm, divided by the extinction coefficient (ɛ) calibrated with standard pigments purchased from DHI Laboratory Products (Denmark). The relative concentration of each pigment was determined by normalizing to the concentration of chlorophyll *a*.

### Thylakoid membrane isolation

Thylakoid membranes were isolated from 6-week-old plants as previously described (Casazza et al., 2001). Briefly, 2 g of leaves were cut into 0.5 cm pieces and incubated in 50 mL of ice-cold water for 20 min. After incubation, the water was drained, and the leaf tissue was blotted dry. Leaf tissue was then ground in 15 mL of grinding buffer [0.4 M sorbitol, 5 mM EDTA, 5 mM EGTA, 5 mM MgCl_2_, 10 mM NaHCO_3_, 20 mM Tricine-NaOH (pH 8.4), 0.5% (w/v) BSA] using a Polytron homogenizer (IKA), gradually increasing the speed to maximum until the sample was fully homogenized. The homogenate was filtered through a Steriflip vacuum filter (20 µm pore size) (Millipore) and centrifuged at 2,600 × *g* for 3 min at 4 °C. The resulting pellet was resuspended with 5 mL of resuspension buffer [0.3 M sorbitol, 2.5 mM EDTA, 5 mM MgCl_2_, 10 mM NaHCO_3_, 20 mM HEPES-NaOH (pH 7.6), 0.5% (w/v) BSA] and centrifuged at 2,600 × *g* for 3 min at 4 °C. The pellet was resuspended with 10 mL of hypotonic buffer [2.5 mM EDTA, 5 mM MgCl_2_, 10 mM NaHCO_3_, 20 mM HEPES-NaOH (pH 7.6), 0.5% (w/v) BSA] and centrifuged at 2,600 × *g* for 3 min at 4 °C. The pelleted thylakoid membrane was resuspended in a small volume of resuspension buffer. Chlorophyll concentration was determined using 80% (v/v) acetone according to Porra et al. (1989).

### SDS-PAGE and immunoblot analysis

Thylakoids were solubilized with 92.5 mM Tris-HCl (pH 8.0), 2.5% (w/v) lithium dodecyl sulfate, and 62.5% DTT for 30 min at room temperature. Non-solubilized fractions were removed by centrifugation at 20,800 × *g* for 5 min at 23 °C. Proteins were separated by SDS-PAGE using TGX Any kD Precast gels (Bio-Rad), and the separated proteins were transferred to a PVDF membrane using a Trans-Blot Turbo transfer system (Bio-Rad). The membrane was then incubated in EveryBlot Blocking Buffer (Bio-Rad) for 10 min. The membrane was washed with TBS-T solution [0.3% (w/v) Tris base, 0.8% (w/v) NaCl, 0.02% (w/v) KCl, 0.1% (w/v) Tween-20, pH 7.4]. Protein abundance was determined using an anti-PsbS antibody (Li et al., 2000). The enhanced chemiluminescence signals, generated by horseradish peroxidase activity on a donkey anti-rabbit secondary antibody (Cytiva), were detected and quantified using a ChemiDoc MP Imaging System (Bio-Rad).

## AUTHOR CONTRIBUTIONS

M.I. and K.K.N. designed the research; R.P. and E.L. performed experiments; R.P., M.I., and K.K.N. analyzed data; M.I. and K.K.N. provided resources and supervision; M.I. prepared figures and wrote the paper; All authors read and approved the contents of this article.

## ACKNOWLEDGMENTS

We thank Dr. Luca Dall’Osto for providing *Arabidopsis* seeds for *npq4-E122Q-E226Q*, *npq4-E122Q*, *npq4-E226Q*, and *koLhcb4*. We also thank Dr. Dhruv Patel-Tupper for technical advice on *Arabidopsis* crossing and guidance during the initial stages of this study. This work was supported by the US Department of Energy Office of Science, through the Photosynthetic Systems program in the Office of Basic Energy Sciences. R.P. acknowledges support from the NIH NIGMS #5T32GM132022 genetics training grant and the Daniel Arnon Graduate Fellowship. K.K.N. is an investigator of the Howard Hughes Medical Institute.

## CONFLICTS OF INTEREST

The authors declare no conflicts of interest.

## Notes

### Competing Interest Statement

The authors have declared no competing interest.

