## Supplementary Information for "Zeaxanthin-dependent rapid energy quenching uncovers complexity of PsbS protonation"

### SUPPLEMENTARY MATERIAL

**Supplementary Table S1.** Primers used for genotyping *LHCB4* genes (de Bianchi et al., 2011).

| Gene | | Primer sequence |
|---|---|---|
| *LHCB4.1* | Forward | 5’-TCACCAGATAACGCAGAGTTTAATAG-3’ |
| | Reverse | 5’-CACATGATAATGATTTTAAGATGAGGAG-3’ |
| | T-DNA | 5’-CATCATACTCATTGCTGATCCATG-3’ |
| *LHCB4.2* | Forward | 5’-TGTCTTGGTCTATTGTACTGTCTCG-3’ |
| | Reverse | 5’-ATATCAATGTCGAGATAGAGAACGG-3’ |
| | T-DNA | 5’-GCTTCCTATTATATCTTCCCAAATTACCAATACA-3’ |
| *LHCB4.3* | Forward | 5’-GTGAGCTGATCCATGGAAGGTGG-3’ |
| | Reverse | 5’-GGCCGGTTTTGAACGATTGATGTGAC-3’ |
| | T-DNA | 5’-GCGTGGACCGCTTGCTGCAACT-3’ |

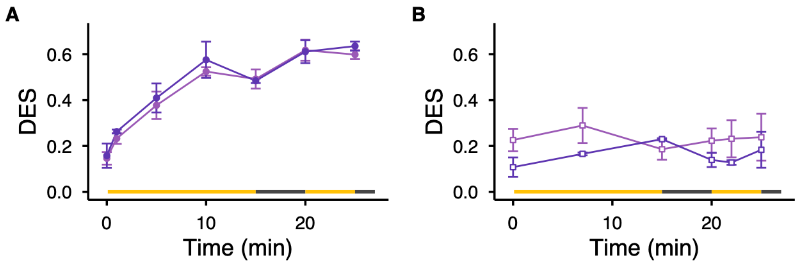


**Supplementary Figure S1. Changes in DES of the single-protonation-site mutants (*E122Q* and *E226Q*) during NPQ measurements. A)** DES of leaf discs was measured by HPLC pigment analysis during NPQ induction and relaxation through two consecutive cycles of AL illumination and dark in *E122Q* (light purple) and *E226Q* (dark purple), as shown in Figure 6. **B)** The DES was measured after 30 min of DTT treatment at 5 mM concentration (open square), as shown in Figure 6. The yellow and black bars indicate the durations of the AL and dark periods. The intensity of AL was ~1000 µmol photons $m^{-2}$ $s^{-1}$. All data points are reported as means ± SD (n = 3).

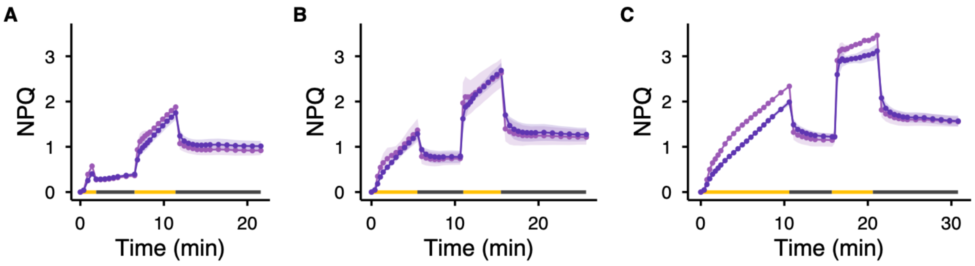


**Supplementary Figure S2. Effect of the initial AL duration on *EQ* jump induction during two light/dark cycles in NPQ measurements for the single-protonation-site mutants (*E122Q* and *E226Q*).** NPQ induction and relaxation kinetics were analyzed in *E122Q* (light purple) and *E226Q* (dark purple). The duration of the first AL was varied: **A)** 1 min, **B)** 5 min, and **C)** 10 min. The yellow and black bars indicate the durations of the AL and dark periods. The intensity of AL was ~1000 µmol photons $m^{-2}$ $s^{-1}$. All data points are reported as means ± SD (n = 3).

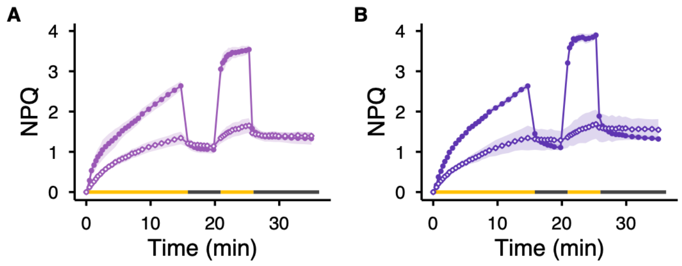


**Supplementary Figure S3. Effect of nigericin treatment on NPQ kinetics for the single-protonation-site mutants (*E122Q* and *E226Q*).** NPQ induction and relaxation kinetics over two consecutive light/dark cycles were analyzed for control measurements (closed circle) in **A)** *E122Q* (light purple) and **B)** *E226Q* (dark purple). NPQ kinetics were measured after 30 min of nigericin treatment at 50 μM (open diamond). The intensity of AL was ~1000 μmol photons $m^{-2}$ $s^{-1}$. The yellow and black bars indicate the durations of the AL and dark periods. All data points are reported as means ± SD (n = 3).

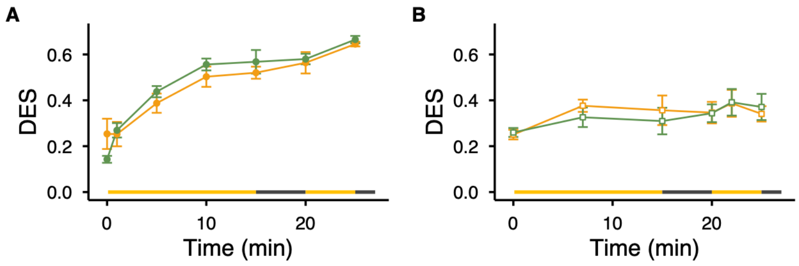


**Supplementary Figure S4. Changes in DES of *koLhcb4* and *E122Q-E226Q koLhcb4* mutants during NPQ measurements. A)** DES of leaf discs was measured by HPLC pigment analysis during NPQ induction and relaxation through two consecutive cycles of AL illumination and dark in *koLhcb4* (orange) and *E122Q-E226Q koLhcb4* mutants (sage green), as shown in Figure 7A. **B)** The DES was measured after 30 min of DTT treatment at 5 mM concentration (open square). The yellow and black bars indicate the durations of the AL and dark periods. The intensity of AL was ~1000 µmol photons $m^{-2}$ $s^{-1}$. All data points are reported as means ± SD (n = 3).

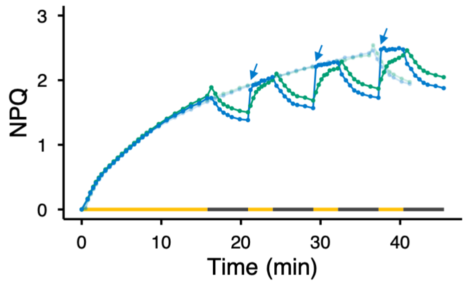


942 
943 **Supplementary Figure S5. NPQ induction and relaxation kinetics shown in Figs. 5A and 5B,**
944 **merged for direct comparison.** Data for *npq4* (green) and *E122Q-E226Q* (blue) are shown; WT
945 is omitted for clarity. Data from Fig. 5A are plotted in a lighter shade to facilitate comparison with
946 Fig. 5B. Arrows indicate where the *EQ* jump reaches the NPQ level observed under continuous
947 AL.
